# Accurate loop calling for 3D genomic data with cLoops

**DOI:** 10.1101/465849

**Authors:** Yaqiang Cao, Xingwei Chen, Daosheng Ai, Zhaoxiong Chen, Guoyu Chen, Joseph McDermott, Yi Huang, Jing-Dong J. Han

## Abstract

Sequencing-based 3D genome mapping technologies can identify loops formed by interactions between regulatory elements hundreds of kilobases apart. Existing loop-calling tools are mostly restricted to a single data type, with accuracy dependent on a pre-defined resolution contact matrix or called peaks, and can have prohibitive hardware costs. Here we introduce cLoops (‘see loops’) to address these limitations. cLoops is based on the clustering algorithm cDBSCAN that directly analyzes the paired-end tags (PETs) to find candidate loops and uses a permuted local background to estimate statistical significance. These two data-type-independent processes enable loops to be reliably identified for both sharp and broad peak data, including but not limited to ChIA-PET, Hi-C, HiChIP and Trac-looping data. Loops identified by cLoops showed much less distance-dependent bias and higher enrichment relative to local regions than existing tools. Altogether, cLoops improves accuracy of detecting of 3D-genomic loops from sequencing data, is versatile, flexible, efficient, and has modest hardware requirements, and is freely available at: https://github.com/YaqiangCao/cLoops.

## Introduction

Three-dimensional genomic interactions are essential for genome organization which provides vital biological function. A loop is classified as two genomic loci that are linearly distant, but have a significantly higher contact frequency than random noise (Yu and Ren 2017). So far, the CTCF (Splinter et al. 2006; Handoko et al. 2011), cohesin (Kagey et al. 2010; Rao et al. 2017), and YY1 (Weintraub et al. 2017) proteins are thought to anchor most of the chromatin loops. These chromatin loops may reveal the transcriptional regulatory roles of distal regulatory elements, such as in enhancer-promoter looping.

Chromatin loops can be identified at near-kilobase resolution (Yu and Ren 2017). With the development of high-resolution chromosome conformation capture (3C) derived high-throughput sequencing methods, it is possible to identify loops genome-wide (Dekker 2016). ChIA-PET (Fullwood et al. 2009; Tang et al. 2015) identifies high-resolution interactions between regulatory elements using target antibodies. Hi-C (Lieberman-Aiden et al. 2009; Rao et al. 2014) maps all possible genomic interactions in an unbiased manner. With deep sequencing (i.e., 6.5 billion total paired-end tags (PETs)), *in situ* Hi-C can achieve 1kb level resolution (Rao et al. 2014), which enables the high-resolution detection of loops. Meanwhile, HiChIP (Mumbach et al. 2016) - combining the advantages of ChIP and *in situ* Hi-C - uses fewer numbers of input cells than ChIA-PET, and attains higher signal-to-background enrichment than *in situ* Hi-C to provide high resolution loops. A new method to identify short and long range interactions called Trac-looping (Lai et al. 2018) was developed recently that uses transposon linkers prior to fragmentation and ligation. Each different technology generates huge datasets and has major computational demands, creating a need for efficient and versatile analysis tools.

Finding long-range loops from 3D genomic interaction data is a computational task equivalent to finding peaks from ChIP-seq data, and is the basic analysis step prior to biological interpretation. Due to the data-type specific technology biases and different resolutions between them, many tools have been designed to call loops. With Hi-C, no algorithm is yet considered to be a golden standard (Forcato et al. 2017). Recently developed loop calling tools for ChIA-PET data such as Mango (Phanstiel et al. 2015) and MICC (He et al. 2015) - implemented in ChIA-PET2 (Li et al. 2016) - often start with peak calling, and then use exhaustive combinations of peaks to find candidate loops, including modeling the relation between paired-end tags (PETs) and distances, and the peaks’ size and depth, which altogether increases data processing time. Importantly, uncertainty in analysis arises when modeling the PETs and distance relations, as different fitting functions and parameters can lead to different loop identification. There is also a problem of bias if the interacting loci forming loops may exist outside of peak regions, which would bias the background used in significance estimations. Correspondingly, we have noticed these tools fail to call loops accurately for data containing broad peaks, such as H3K4me1 ChIA-PET data. The hardware requirements for loop calling from Hi-C data present another major limitation. For example, the major Hi-C loop-calling tool HiCCUPS from Juicer (Durand et al. 2016b) requires NVIDIA GPUs, which are more expensive than random access memory (RAM) (e.g., a TITAN Xp is about 10-fold more expensive than 16GB RAM) and may be incompatible with many previous server setups. Due to huge PET numbers, loop-calling tools for Hi-C usually have high RAM usage. However, according to estimates in a Hi-C tools comparative study, contact matrix based tools like Fit-Hi-C (Ay et al. 2014) and GOTHiC (Mifsud et al. 2017) require more than 512 GB of RAM for a 5kb resolution contact matrix (Forcato et al. 2017), making the loops calling on a 1kb high-resolution contact matrix from deep sequencing impossible. Currently, to our knowledge, there is only one targeted loop calling tool for HiChIP data, hichipper (Lareau and Aryee 2018). The basic loop calling procedure of hichipper is very similar to Mango and ChIA-PET2; it first uses MACS to call peaks from the HiChIP data with custom background models and then depends on Mango to identify loops. The method for calling loops from Trac-looping data the same as in hichipper, thus the biases can also be inherited from those in Mango.

To avoid biases present in the existing loop calling tools and to enable a low-computational-cost and universal method for 3D-genome mapping data we developed a new tool: cLoops (“see loops”). cLoops is a versatile loop calling tool for multiple 3D-genome mapping data. It uses an unbiased clustering algorithm to find candidate loops, coupled with a permutated local background method for estimation of a candidate loop’s statistical significance. We show the advantages of cLoops over existing state-of-the-art loop calling tools by comparisons with ChIA-PET, Hi-C, HiChIP and Trac-looping data. Briefly, 1) cLoops is easy to use, having only two essential input parameters, for which we provide predetermined default values for ChIA-PET, Hi-C, HiChIP and Trac-looping data. 2) cLoops can run efficiently on PCs and accurately identify loops for both sharp-peak and broad-peak data. 3) Compared to other tools, performance was distinguished by cLoops’ uniquely identified loops that showed more easily distinguishable signals within their neighboring regions, cLoops identified more distant loops from Hi-C and HiChIP data, and showed higher overlap with ChIA-PET loops. 4) cLoops’ reliability was not affected by sequencing depth, with equivalent performance in both deep and unsaturated HiChIP sequencing data. 5) cLoops is not tied to any particularly experimental method therefore is applicable to 3D-genome mapping data generated by future experimental methods, as long as there are data with enriched interactions detectable on an interaction heatmap.

## Results

### The cDBSCAN algorithm

DBSCAN (Density-Based Spatial Clustering of Applications with Noise) (Ester et al. 1996) is one of the most widely used unsupervised clustering algorithms. DBSCAN contains two key parameters: *eps* defines the distance within which for two points are classified as neighbors and *minPts* defines the smallest number of points required in a cluster. It has been introduced for ChIA-PET by taking all paired-end tags (PETs) as 2D points, and identifying significant clusters in 2D space as potential loops (Chepelev et al. 2012). The density-based principle, tolerance of noise, and unsupervised auto-determination of the number of clusters, theoretically make DBSCAN very suitable for finding candidate loops from 3D genomic interaction data. However, the original DBSCAN algorithm runs very slowly for ChIA-PET and Hi-C data (with complexity of *O* (*N^2^*) without any optimization for neighbor search, *N* is the number of points). For example, if implemented with the C programming language based KD-Tree for neighbor search (named kDBSCAN, with complexity of *O* (*Nlog(N))*, **Methods**) on a computer with a 3.2G CPU (see the detailed configuration of computers used in **Supplemental Information**), the average time of 5 runs for kDBSCAN is about 32 seconds (*eps* = 5, *minPts* = 750) to finish clustering on 99,674 PETs in the smallest human autosome (chromosome 21) from GM12878 CTCF ChIA-PET data (**Supplemental Table 1**) and about 1.1 hours (*eps* = 20, *minPts* = 5000) for 2,268,476 PETs in chromosome 21 from GM12878 Hi-C data (**Supplemental Table 1**). Although DBSCAN has been implemented in TADLib for interaction block analysis within topological associated domains (TADs) (Wang et al. 2015), so far no tools have implemented it for loop calling, or to determine the loop calling effectiveness or significance. We thus first propose a specific improvement to DBSCAN (named cDBSCAN for cLoops’ DBSCAN) for 2D data, by introducing an indexing method for noise reduction and neighbor search (see toy example data in **Figure 1A** for illustration of the algorithm).

**Figure 1.**
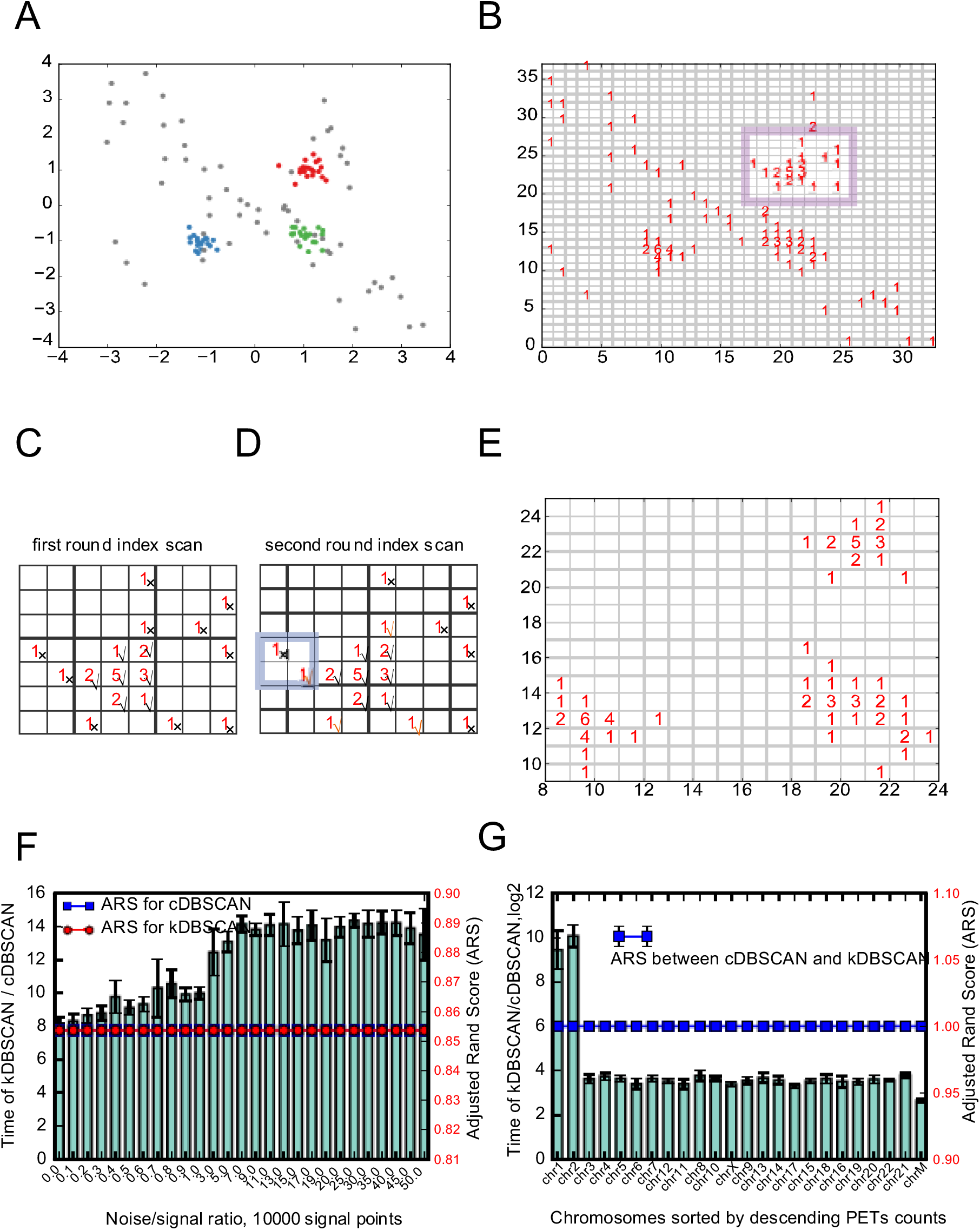
The cDBSCAN algorithm. (A) A toy example of the simulated test data, mainly 3 clusters, centered at (-1,-1), (1,-1) and (1, 1) with *std* = 0.2, a total of 60 signal points and 60 noise points. Noise is generated randomly and marked as grey points. (B) Indexing result, each point is attached to a square with side length of *eps*, which equals to*std* here, the numbers in the squares indicate the number of points indexed in that square. The highlighted region is used to represent detected noise. (C) An example of first round of noise removal, the region is highlighted in (B). For an *eps* square, scan the nearby 8 squares, if the total number of nearby points is less than required *minPts* which is 5 here, then the index square is marked as noise. A noise index is marked by a cross, while a signal index is marked by a checkmark. (D) Second round of noise removal for the same region in (C), for a noise index detected in (C), if one of its neighbor square index is a signal index, then it is re-marked to a signal index. Examples are marked by orange checkmarks. The highlighted region is an example and an outer index that is not re-marked as signal index. (E) An example of indexed *eps* square after noise removal. (F) Comparison of running CPU time at different noise/signal ratio based on 10 repeats for the simulation data. Left y-axis marks the bars for running time ratios; right y-axis marks the lines for adjusted rand scores (ARS). The two ARS are exactly the same. (G) Comparison of running CPU time using real GM12878 CTCF ChIA-PET data (GSM1872886) for each chromosome based on 5 repeats, with *eps* = 750 and *minPts* = 5. Error bars denote standard deviations.

cDBSCAN also has two key parameters with the same meaning as those in DBSCAN: *eps* and *minPts*. For 3C-based genome wide sequencing data like ChIA-PET, HiChIP and Hi-C data, the Manhattan distance (also known as city block distance) is suitable for measuring the absolute position difference for two PETs. Unless specifically mentioned, the distance measurements hereafter refer to Manhattan distance. For the 2D dataset *D* (*X*, *Y*) in the cDBSCAN algorithm, X and Y can be integer or float values, but for loop calling, they are all both integers corresponding to genomic coordinates. We mark the minimum *X*, *Y* as *minX*, *min Y* for the 2D space. cDBSCAN indexes each point (*X*_i_, *Y*_i_) as 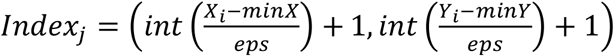 which means each point is assigned to a square whose side length is *eps* (0.2 is used for the toy example) (**Figure 1B**) and *j* marks the index id of the point id *i*. For an indexed *eps* square, if the square together with its surrounding 8 squares contains points fewer than *minPts* (5 is used for the toy example), then it is defined as a noise index. We highlight a region in Figure 1B (also used in Figure 1C and D) to show how cDBSCAN removes noise. There are two rounds of index scanning in cDBSCAN to detect noise. The first round finds all potential noise indexes (marked by a cross in **Figure 1C**), and the second round only searches previously detected noise indexes (cross marked indexes). If there is a first round signal index (marked by checkmarks) in any of its 8 neighbors, then it is marked as a signal index (**Figure 1D** orange checkmarks). The highlighted region in **Figure 1D** is an example showing an outer index that is not re-marked is a signal index and a closer index that is re-marked as a signal index. The idea benefits from k-Nearest Neighbor (KNN) algorithm - that is, if all neighbors are noise then the index is noise. A signal index detected in the second-round search is not counted as a signal index when a noise is corrected back to a signal (**Figure 1D**). This indexing process reduces the search space and saves memory (**Figure 1E**). After indexing, the clustering is performed the same as DBSCAN for the remaining points, but uses the 3×3*eps* squares for neighbor search.

We first evaluated the performance of cDBSCAN by comparing to a C coded KD-tree for neighbor search (termed it kDBSCAN as mentioned above) (**Methods**) using the simulated data. We set 10,000 signal points of 100 clusters and different noise/signal ratios for the simulation data (**Methods**). cDBSCAN coded in pure Python gives the exact same result as kDBSCAN (measured by Adjusted Rand Score (ARS)) (Hubert and Arabie 1985). ARS measures the similarity between clustering results ranging from-1.0 to 1.0, with 0 indicating random labeling and 1 a perfect match. cDBSCAN had reduced memory usage and improved speed (8-16 fold, without considering the inefficiency of Python compared to C for the simulation data (**Figure 1F**). We also validated the speed increase on real GM12878 CTCF ChIA-PET data and found a ~8-1000 fold increases (**Figure 1G**). Comparing to kDBSCAN *O* (*Nlog*(*N*)) complexity, cDBSCAN is *O*(*N*) complexity in most ideal situation, which is further validated by running cDBSCAN for the PETs in chromosome 1 for CTCF ChIA-PET (**Supplemental Figure 1A**), GM12878 Hi-C (**Supplemental Figure 1B**), GM12878 cohesin HiChIP (**Supplemental Figure 1C**) and the Trac-looping data (**Supplemental Figure 1D**) as the run time increases nearly linearly as the PETs increase.

### The overview of cLoops

Based on cDBSCAN we built cLoops (see loops) as a two-step loop-calling algorithm. cLoops is a light-weight tool coded in pure Python with dependence on only a few commonly used and well maintained packages such as scipy, numpy, pandas, joblib and seaborn. The first step uses cDBSCAN to find candidate loops from mapped PET data, without binning PETs into a specific assigned resolution contact matrix (as usually occurs with Hi-C loop-calling tools such as Fit-Hi-C) and without first identifying peaks from PETs and then finding the significant combinations of peaks (as needed by common ChIA-PET loop-calling tools such as Mango). In the second step the estimation of candidate loops’ significance is compared to permuted local backgrounds. The overview of data processing steps of cLoops is demonstrated in **Supplemental Figure 1E**. We show the algorithm details of cLoops using the GM12878 CTCF ChIA-PET data as follows.

First, each intra-chromosomal PET is mapped to a 2D space by taking the middle coordinate of the left-end tag as the X-coordinate, and the middle coordinate of the right-end tag as Y-coordinate into *X_i_*, *Y_i_* where *i* indicates the PET id (**Figure 2A**). All PETs are therefore clustered by cDBSCAN. After clustering, each cluster can be marked as ((*X_K,min_*, *X_K,max_*), (*Y_K,min_*, *Y_K,max_*)), where *k* is the cluster id, *X_K,min_* is the left boundary of left anchor (*X_K,min_* equals *x*_1_ in **Figure 2A**), *X_K,max_* is the right boundary of left anchor (*X*_*K,max*_equals *x*_2_ in **Figure 2A**), *Y_K,min_* is the left boundary of right anchor (*Y_K,min_* equals *y*_1_in **Figure 2A**), *Y_K,max_* is the right boundary of left anchor (*Y_K,max_* equals *y*_2_ in **Figure 2A**). A model based distance cutoff was used to filter out potential self-ligated PETs (**Figure 2B**) (**Methods**). If there are fewer PETs in the inter-ligation clusters than *minPts*, such clusters are removed. The remaining inter-ligation clusters are treated as candidate loops which then have their significance estimated against the local background.

**Figure 2.**
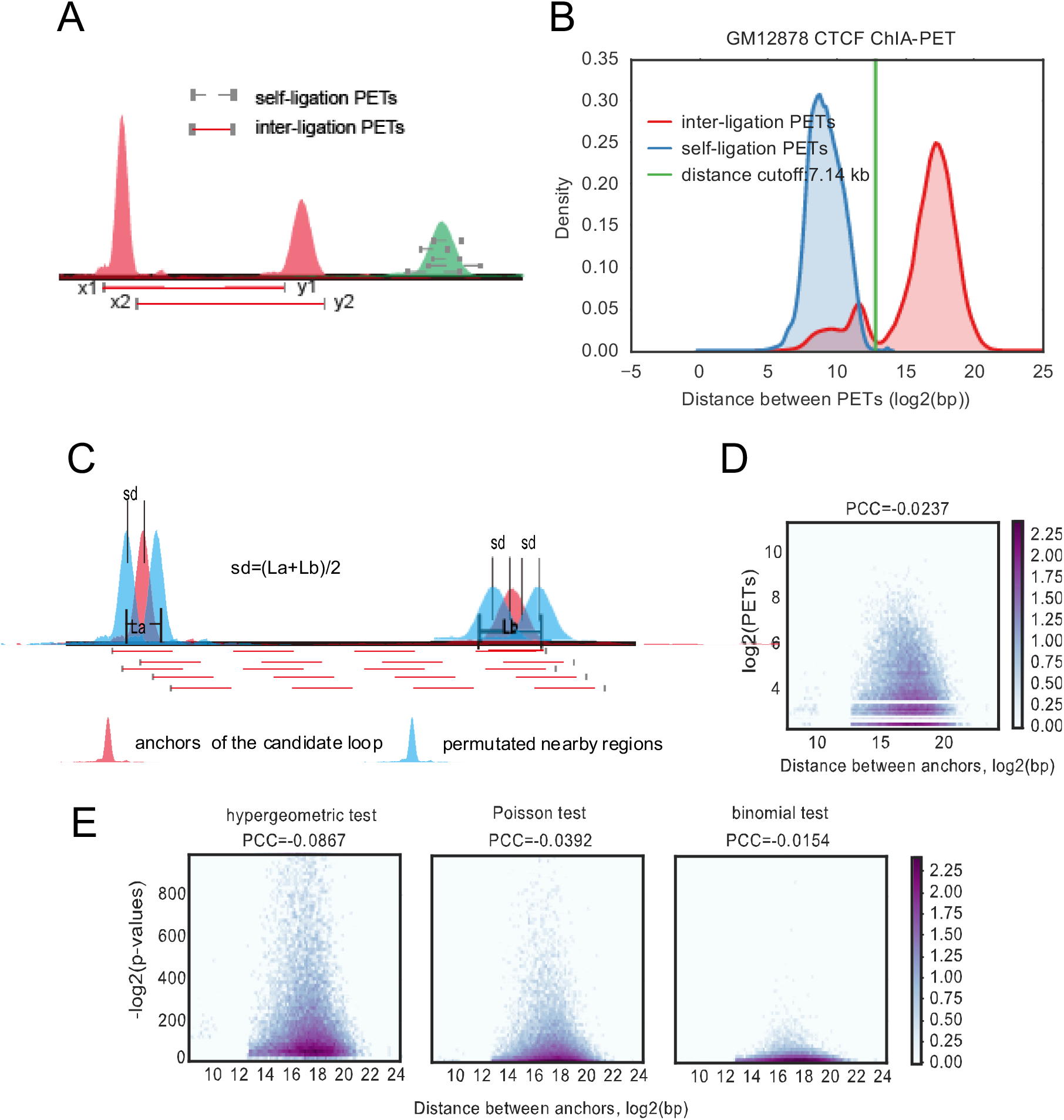
Overview of cLoops. (A) To carry out clustering, each PET is mapped to 2D space as its middle coordinate of left PET mapped to X-axis and right mapped to Y-axis. (B) Distance distribution for GM12878 CTCF ChIA-PET PETs in classified inter-ligation and self-ligation clusters (**Methods**). (C) Permutated local background for estimating candidate loops statistical significance. For two anchors of a candidate loop, all combinations of their upstream and downstream 5 moving windows with size of anchors and step size of the mean length of these two anchors. The mean distance for all combinations is exactly the same as the interacting loop region. (D) Hexbin plot of detected PETs and distance between loop anchors for CTCF ChIA-PET data. (E) Hexbin plot of estimated p-values using different methods and the distance between loop anchors for CTCF ChIA-PET data.

The key parameters used in cLoops are those used to run cDBSCAN, *eps* and *minPts*. *minPts* determines the least number of PETs required for a loop, and *eps* defines the distance for two PETs to be neighbors and this setting is more data dependent. Multiple *eps* and *minPts* can be assigned to cLoops to run cDBSCAN clustering multiple times to find merged consensus candidate loops. Empirically determined parameters were used for ChIA-PET, HiChIP, Hi-C and Trac-looping data (**Methods**).

### Permuted local background for estimating significance of candidate loops

For the second step, to test the significance of a candidate loop over the nearby genomic background, a permuted local background (PLB) is used (**Figure 2C**). Linearly closer anchors in the genome have higher probabilities to capture more PETs linking them due to experimental ligation bias in both ChIA-PET and Hi-C (Paulsen et al. 2014), which needs to be modeled and corrected for in loop significance tests. We designed this permuted local background to save the effort of correcting PET distance bias. For each candidate loop (red peaks), PLBs are defined as all combinations of their upstream and downstream 5 moving windows (light blue peaks, one upstream and one downstream PLB plotted for the left/right anchor, in cLoops 5 moving windows are used to obtain 100 permuted background regions) with the same length as the loop anchors (**Figure 2C**). The shifting size for the moving windows is the mean length of these two anchors. Thus, the mean distance of all permutated windows is exactly the same as the candidate loop. Based on the PLB, the commonly used hypergeometric test, Poisson test and binomial test were used together to determine a candidate loop’s statistical significance; the details of the mathematical model and cutoff are described in **Methods**.

For cLoops-called loops, due to the density-based clustering method and removal of suspected self-ligation PETs based on distance distributions, PET numbers are actually independent of loop distances. For example, in the CTCF ChIA-PET data, the Pearson correlation coefficient (PCC) is −0.0237 between PETs numbers and distances between anchors (**Figure 2D**). The p-values derived using different statistical tests are also independent of loop distances (**Figure 2E**).

### cLoops application to ChIA-PET data

We compared cLoops with three peak-calling based tools loop-calling tool, ChiaSig (Paulsen et al. 2014), ChIA-PET2 (Li et al. 2016) and Mango (Phanstiel et al. 2015) (**Supplemental Table 2**) using multiple ChIA-PET datasets (**Supplemental Table 1**). These three ChIA-PET tools were selected because they are the most frequently used. The run time for these tools is shown in **Supplemental Table 3**. cLoops is designed with parallel computing, while other tools were not, however, even when cLoops was run with only one CPU it was still much faster than ChiaSig and ChIA-PET2 on the GM12878 CTCF and RAD21 ChIA-PET, the HeLa CTCF ChIA-PET data, and K562 H3K4me1 ChIA-PET data (**Supplemental Table 3**).

Heatmaps and global quality of loops were visualized with mean profile heatmaps of loops (centerNormedAPA heatmaps) and the mean P2M (Peak to Mean) scores, respectively (**Methods**). The centerNormedAPA heatmaps were generated by Juicer APA. In a centerNormedAPA heatmap, loops are aligned in the center, and a high contrast ratio compared to the nearby regions indicates good loop quality. If there are highly interacting regions other than the center in a centerNormedAPA heatmap, it indicates either that there are shifts of loop boundaries or global loop quality is not good. As a quantitative indicator for enrichment of loops compared to nearby regions, P2M (computed by Juicer APA), is defined as the ratio of the central pixel to the mean of the remaining pixels (Durand et al. 2016b). In addition to P2M scores, we also show the global mean P2LL scores (Peak to Lower Left) and the related ZscoreLL scores (suggested by the Juicer documentation) for comparison (**Supplemental Figure 7)**. A comparison of the loop anchor size distributions indicates that cLoops can identify loops with a larger range in anchor size than the other peak identification based algorithms, some of which have a predefined anchor size (**Supplemental Figure 8**).

In general, cLoops and Mango outperformed ChiaSig and ChIA-PET2 for all tested ChIA-PET data as indicated by the mean profile heatmaps and the mean P2M scores for ChIA-PET data contain sharp peaks (e.g. CTCF and RAD21) (**Supplemental Figure 2**). We noticed that Mango, ChiaSig and ChIA-PET do not work well with histone modification ChIA-PET data, such as with K562 H3K27ac and H3K4me1 datasets. Mango, ChiaSig and ChIA-PET2 identified limited loop numbers, and the loops’ qualities were worse compared to cLoops, as evaluated both by mean profile heatmaps of loops and the mean P2M scores (**Figure 3**A, **Supplemental Figure 2 and Supplemental Figure 3B).** The cumulative aggregate peak analysis (CAPA) designed by Mango to evaluate quality of loops called from ChIA-PET data through Hi-C data was used to further compare performance. CAPA validated advantages of cLoops and Mango over ChiaSig and ChIA-PET2 (**Supplemental Figure 3A**) in enriching for Hi-C interacting signals for ChIA-PET data containing broad peaks (**Figure 3B**), and similar performances of cLoops and Mango for ChIA-PET data containing sharp peaks (**Supplemental Figure 3B**). Worse performances are partially due to use of the narrow peak calling model of MACS (Zhang et al. 2008) as a default setting. We show two randomly selected unique loops called by cLoops from H3K4me1 ChIA-PET data (**Figure 3C**) and H3K27ac ChIA-PET data (**Figure 3D**) as examples to illustrate cLoops’ ability to detect reliable loops that could be observed from the visualization of raw PETs which are missed by other tools. Moreover, Mango estimated p-values showed a higher dependence on the anchors’ distance, showing higher significance for closer anchors, which suggests insufficient correction for the experimental bias (**Supplemental Figure 3C**).

**Figure 3.**
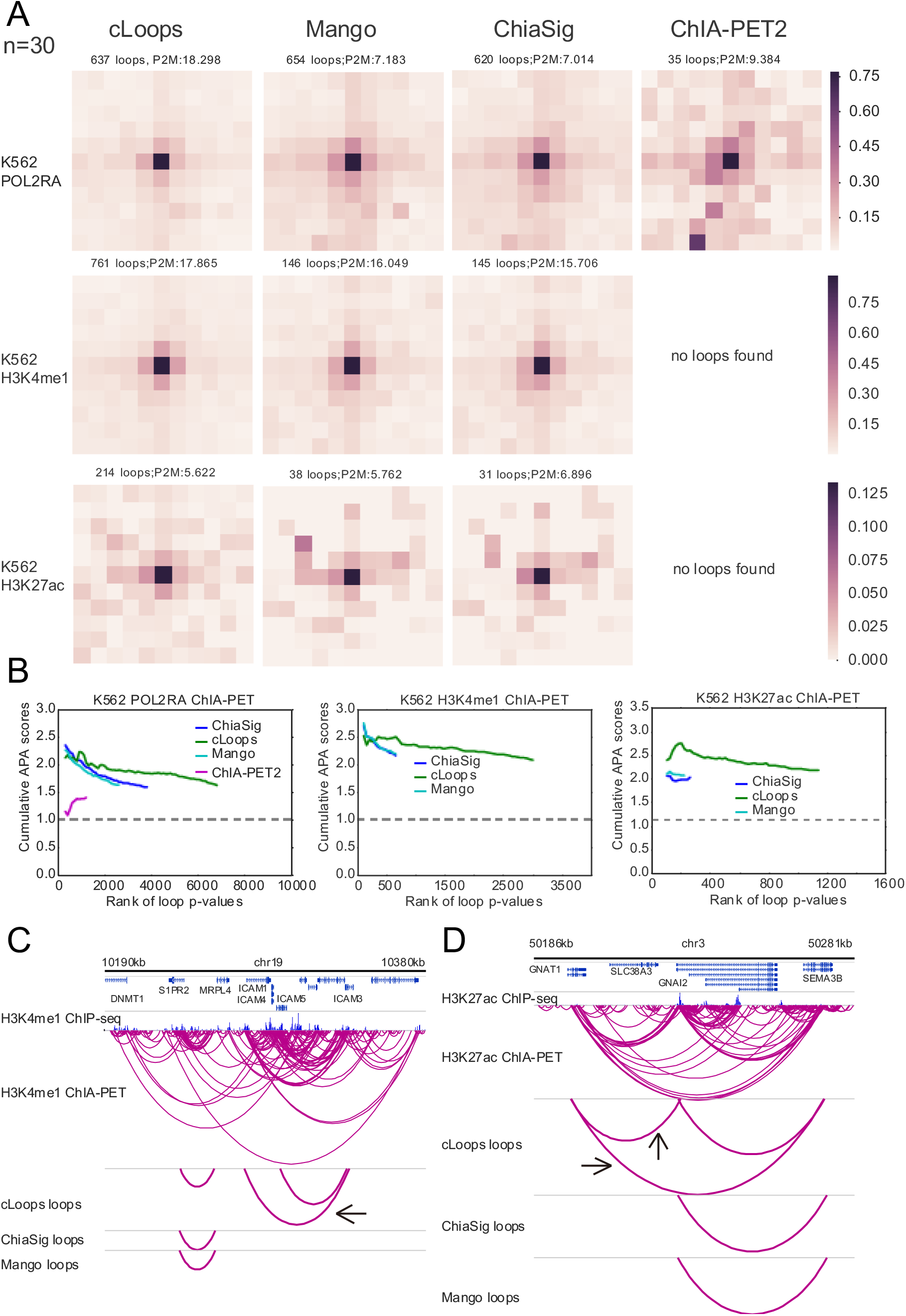
cLoops applied to ChIA-PET data and comparison with other tools. (A) centerNormedAPA heatmaps from Juicer (Durand et al. 2016b) aggregate peak analysis (APA) were shown for loops obtained by cLoops from K562 POLR2A, H3K27ac and H3K4me1 ChIA-PET data. The number of loops and P2M score from whole genome-wide analysis were annotated at head of each dataset heatmap. The P2M score is the mean of all P2M values, which indicate the enrichment of loops compared to nearby regions. In the Juicer APA analysis, -n was set to 30 (default parameter) to analysis loops with anchor distance >= 150kb. More comparisons for distance filtered loops are shown in **Supplemental Figure 2.** (B) Cumulative APA (CAPA) for evaluating the qualities of loops called from ChIA-PET data using Hi-C data. Higher scores mean the loops are better supported by Hi-C (APA score > 1.0). (C, D) Example of unique loops detected by cLoops for H3K4me1 (C) and H3K27ac (D) ChIA-PET data.

### cLoops application to Hi-C data

We compared cLoops with five Hi-C loop calling tools recently evaluated in a tool-performance comparison study (Forcato et al. 2017), namely diffHic (Lun and Smyth 2015), Fit-Hi-C (Ay et al. 2014), GOTHiC (Mifsud et al. 2017), HiCCUPS (Durand et al. 2016b) and HOMER (Heinz et al. 2010) (**Supplemental Table 2**), using the high resolution deep sequencing data from GM12878 and K562 Hi-C data (**Supplemental Table 1**). To compare performance on the same hardware system, we used a PC system (**Supplemental Information**) and equivalent pre-processing with HiC-Pro. We did not compare HIPPIE (Hwang et al. 2015) for following reasons: 1) HIPPIE requires the Sun Grid Engine system and to compare tools based on equivalent systems we could only access a PC system with GPUs. 2) HIPPIE pre-processing differs because it uses STAR (Dobin et al. 2013) for mapping and requires its own pre-processing pipeline. 3) HIPPIE didn’t show unique advantages for calling loops in the comparison study (Forcato et al. 2017). Parameters and loops selections were mostly set according to those used in a previous comparison study (Forcato et al. 2017) (**Supplemental Table 2**). Raw FASTQ data was first processed by HiC-Pro and the required input files for each tool were converted from HiC-Pro output files (**Methods**). The runtimes for these tools are available as **Supplemental Table 4**.

For both GM12878 and K562 Hi-C data, a region on chromosome 21 (36,000 to 39,500kb) contained 6 obvious conserved, visibly salient loops in the 5kb resolution heatmaps (5kb resolution was chosen for visualization in Juicebox to get clear views of loops and 5kb is default most high-resolution setting for a .hic file visualized in Juicebox), designated as “a”, “b”, “c”, “d”, “e”, “f” (note that there are actually 2 loops at the “e” region if further zoomed in) (**Figure 4A and B**). We compared the loops detected by different tools for this example region for both Hi-C and the following HiChIP data. Generally, cLoops and HiCCUPS outperformed other tools in detecting most of the visible loops, while also avoiding detection of probable false-positives located near the heatmap diagonal for both GM12878 and K562 data (**Figure 4A and B**). More examples of visible loop comparisons are shown in **Supplemental Figure 5**. The mean loop profile heatmaps and mean P2M scores indicated that the majority of loops detected by diffHic, Fit-Hi-C, GOTHiC and HOMER are located very near to the diagonal line and have no enriched interaction signals compared to nearby regions. The distribution of distances between loop anchors also supported this conclusion, as HOMER and GOTHiC tended to identify closer loops, thus showing distance dependency (**Supplemental Figure 4E**). The mean profile heatmaps showed cLoops had higher enrichment of interacting signals of loops compared to nearby regions, while loops from HiCCUPS showed no enrichment at the center of called loops (center on heatmaps) compared with its upper left corner region (**Figure 4A and B**).

**Figure 4.**
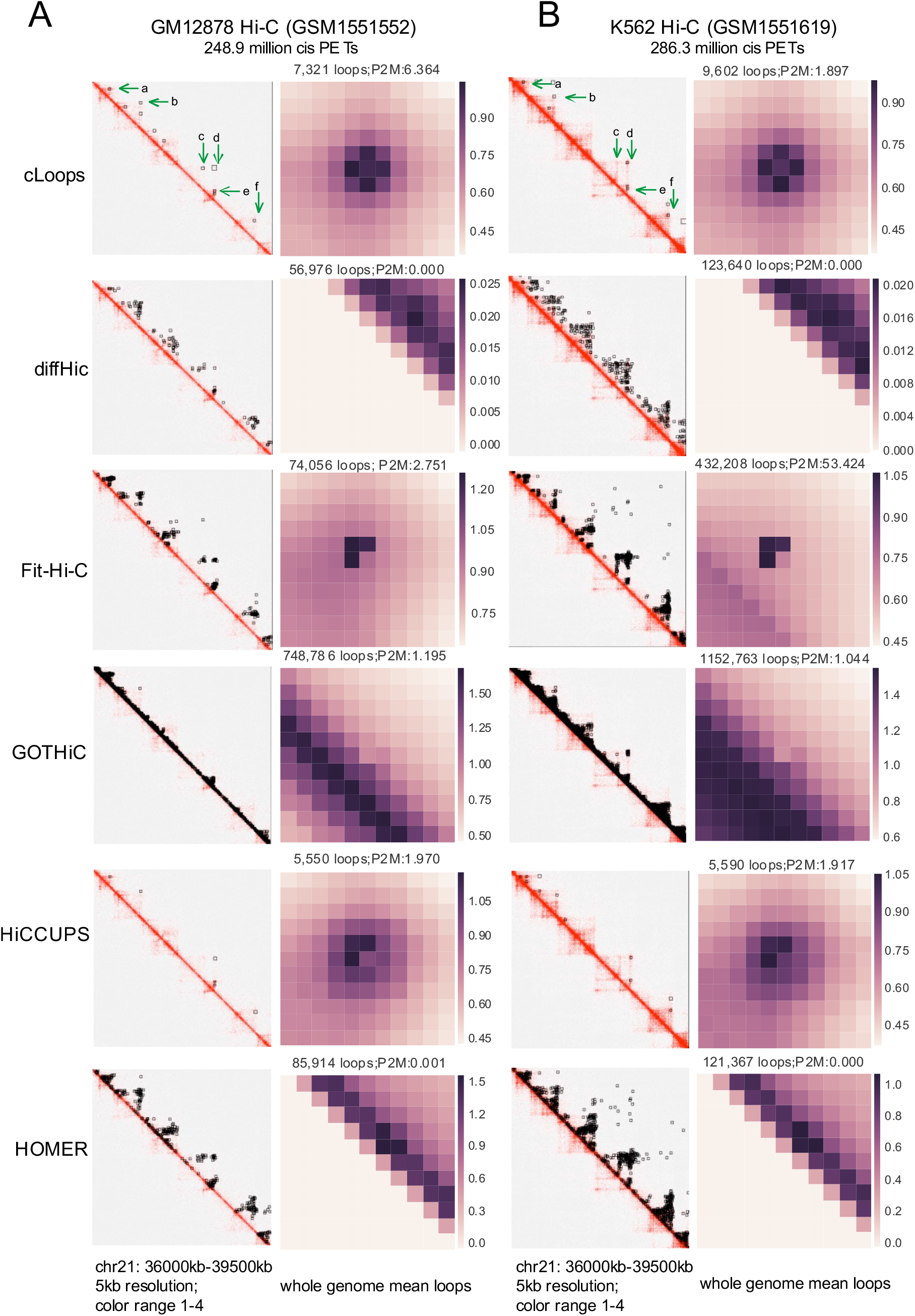
cLoops applied to Hi-C data and comparison with other tools. (A) Example of loops called by each tool for the same region (left) and APA for called loops (right) for GM12878 Hi-C data. centerNormedAPA heatmaps from Juicer APA analysis were shown for the mean loops. The number of loops and P2M score from whole genome-wide analysis were annotated at head of each dataset heatmap. (B)Comparison for K562 Hi-C data.

For independent confirmation, the higher overlap of cLoops and HiCCUPS called GM12878 Hi-C loops with CTCF and RAD21 ChIA-PET loops (**Supplemental Figure 4A and B**) and HiChIP loops (**Supplemental Figure 4C**) supported the robustness of performance of cLoops and HiCCUPS over other tools. That is, the higher mean density of CTCF, RAD21 and SMC3 ChIP-seq binding signals on anchors called by cLoops and HiCCUPS strongly support their higher accuracy and comparatively better performance. Moreover, we also observed two distinct advantages of cLoops compared to all other tools: 1) for called loops, the PET numbers are less dependent on distance between anchors (**Supplemental Figure 4D**), and 2) cLoops can better detect more distant loops (**Supplemental Figure 4E**).

HiCCUPS is mainly based on comparing observed values to expected values for every pixel (where pixel size depends on the pre-defined resolution for contact matrix), and then determining the significance for the pixel using a modified Benjamini-Hochberg FDR control procedure (so called “*λ*-chunking”), with additional filters for local neighborhoods. Then the loops are clustered from the significant pixels. The concept of the HiCCUPS algorithm is quite different from cLoops; the setting of Hi-C specific “*λ*-chunking” and the additional filters may limit HiCCUPS to other 3D-genomic data, and the time-consuming pixel level computing is also limited to an inside loops distance cutoff (<=2MB), while cLoops does not have such limitations. Overall, cLoops’ loops are better supported by ChIA-PET and HiChIP data overlap in GM12878 and showed less bias against distant loops, and cLoops does not need GPUs to run.

### cLoops application to deep-sequencing HiChIP data

Although Fit-Hi-C and Mango were used for calling loops in their original HiChIP method paper (Mumbach et al. 2016), only HiCCUPS called loops using merged PETs from biological and technical replicates were provided as Supplemental data, so we first compared cLoops to HiCCUPS using the merged GM12878 cohesin HiChIP data.

cLoops obtained similar numbers of loops as HiCCUPS for the GM12878 cohesin HiChIP data on the example chromosome 21 region mentioned above in the Hi-C comparisons (**Figure 4**), where cLoops detected all 6 visible loops (**Figure 5A**). HiCCUPS did not detect loop “f” despite detecting the “f” loop in Hi-C data (**Figure 5A**). The mean loops profile heatmaps indicated HiCCUPS may detect more loops close to the heatmap diagonal line (**Figure 5 B and C**). We validated this by showing the distance between anchors for all loops (**Figure 5F**), and with the unique loops mean profile heatmap for cLoops and HiCCUPS (**Figure 5G**) and the distance between anchors for unique loops (**Figure 5I**), which altogether show cLoops can detect more distant loops and the loops with higher signal enrichment. Furthermore, the loops called by cLoops are better supported by both ChIA-PET loops and Hi-C loops for all called loops (**Figure 5E**), and for the unique loops (**Figure 5H**). Moreover, the loop anchors called by cLoops have higher CTCF, RAD21 and SMC3 ChIP-seq tag densities than those of HiCCUPS (**Figure 5D**).

**Figure 5.**
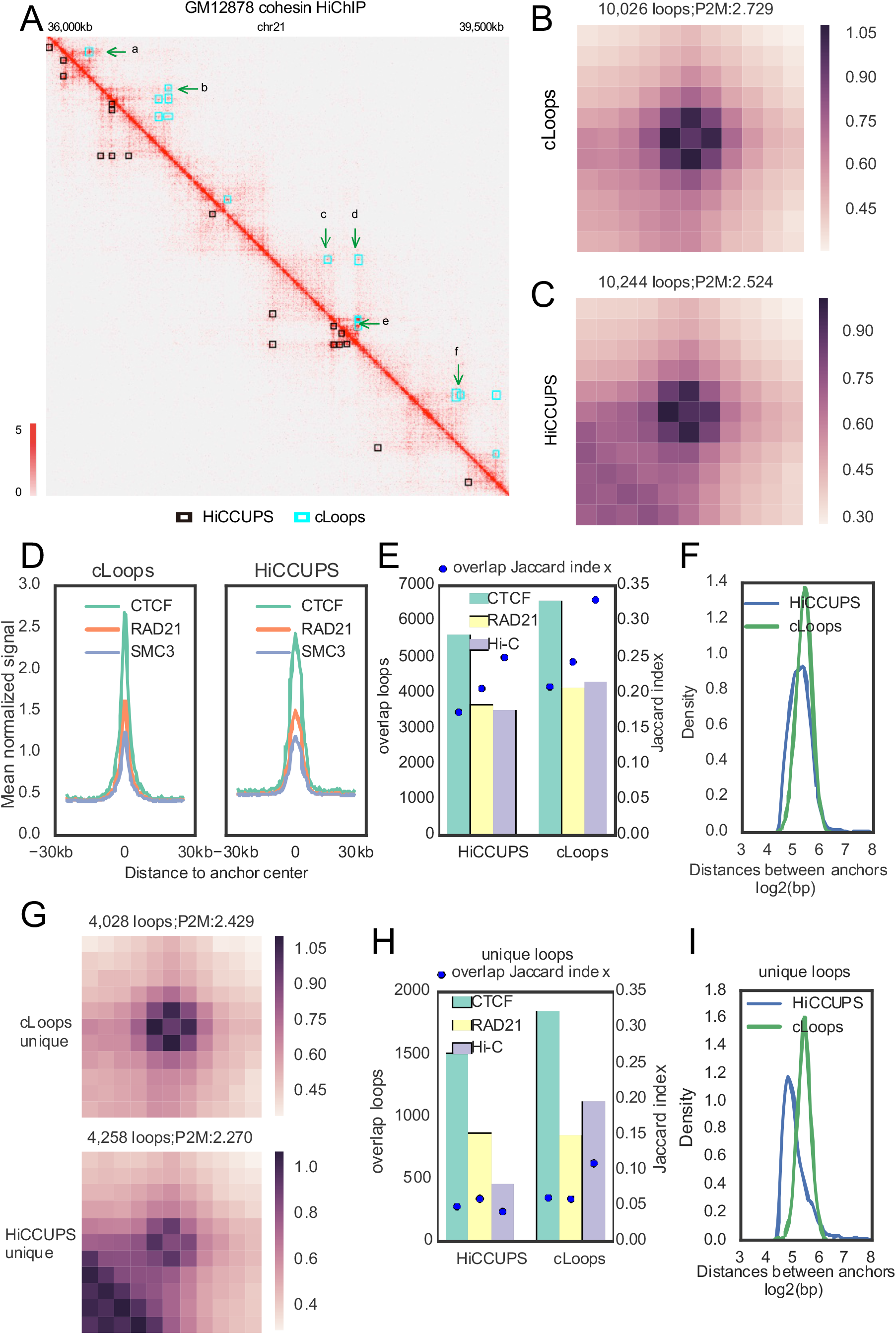
cLoops applied to deep sequenced HiChIP data and comparison with HiCCUPS. (A) Examples of loops called by cLoops (upper) and HiCCUPS (lower) for merged GM12878 cohesin HiChIP data. The loops called by HiCCUPS were obtained from the HiChIP original paper. (B) Mean profile heatmap of loops called cLoops. (C) Mean profile heatmap of loops called by HiCCUPS. (D) Mean ChIP-seq signal of CTCF, RAD21 and SMC3 on loop anchors. (E) Loops overlapping with CTCF ChIA-PET loops, RAD21 ChIA-PET loops and Hi-C loops. (F) Distribution of distances between loop anchors for all loops. (G) Mean profile heatmap of unique loops detected by cLoops (upper) and HiCCUPS (lower). (H) Uniquely detected loops overlapping with CTCF ChIA-PET loops, RAD21 ChIA-PET loops and Hi-C loops. (I) Distribution of distances between loopanchors of uniquely detected loops.

### cLoops application with low-depth sequencing HiChIP data

With the capture enrichment process, HiChIP could in principle reveal enriched loops with under-sequenced PETs compared to Hi-C. Therefore, we wondered whether cLoops’ performance is still relatively good in this situation. We compared cLoops with the Hi-C loop calling tools compared above and hichipper (Lareau and Aryee 2018) (**Supplemental Table 2**) using the two technical replicates of biological replicate 1 from the cohesin GM12878 HiChIP data. The run time for these tools is shown in **Supplemental Table 5**. Even when using only one CPU, cLoops was faster than HOMER, hichipper, Fit-Hi-C and GOTHiC.

The performances of each tool were assessed in a similar way as for Hi-C data above. For the low-depth sequenced HiChIP data, in summary, 1) cLoops, HiCCUPS, HOMER and hichipper can obtain similar visible loops as shown in the example region (**Figure 6A**), detecting majority of the four example loops (“a”, “b”, “c”, “d”) on the heatmaps for both replicates, and not detecting artificial interaction signals close to diagonal line. Also, the mean profile heatmaps for all loops from all four tools showed enrichment over nearby regions, while loops from diffHic and GOTHiC showed obvious patterns close to the diagonal line (**Figure 6B**). 2) Loops called by cLoops, HiCCUPS and HOMER were consistent with CTCF ChIA-PET loops (**Supplemental Figure 6A**), RAD21 ChIA-PET loops (**Supplemental Figure 6B**) and Hi-C loops (**Supplemental Figure 6C**), as measured by the Jaccard index. 3) The detected PET numbers in loops called by cLoops and HiCCUPS are far less dependent on distance between anchors than with other tools (**Supplemental Figure 6D**). The distance dependence is especially high for HOMER, hichipper and Fit-Hi-C. 4) cLoops, HiCCUPS and Fit-Hi-C could detect more distant loops compared to others (**Supplemental Figure 6E**). Notably, cLoops does not need additional control parameters like –L and –U in Fit-Hi-C to detect distant loops. 5) HiCCUPS and cLoops had the highest Jaccard Index of overlapping loops between technical replicates, except for GOTHiC, as it appeared to call too many loops (e.g., the example region called loops at nearly all positions), whereas hichipper showed the lowest Jaccard Index, indicating the peak-based strategy might be biased by errors in peak calling (**Supplemental Figure 6F**). 6) The anchors of loops detected by cLoops, HOMER and hichipper are better supported by the CTCF, RAD21 and SMC3 ChIP-seq data (**Supplemental Figure 6G**). Due to the first pre-customized peak-calling step of hichipper, the higher enrichment of ChIP signal on hichipper anchors is expected by design. 7) Again, cLoops does not need GPU like HiCCUPS.

**Figure 6.**
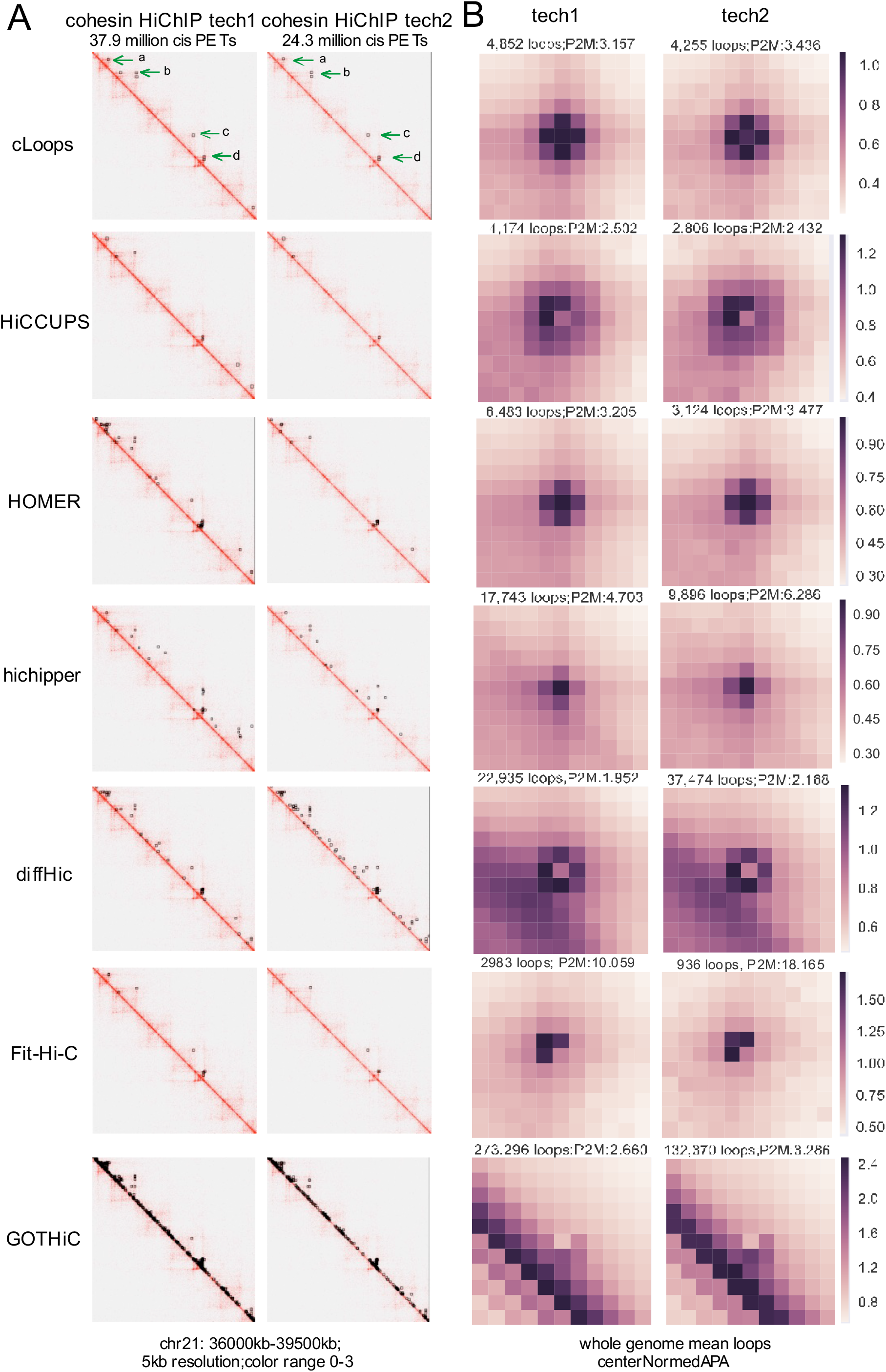
cLoops applied to under-sequenced HiChIP data and comparison with other tools. (A) Examples of loops called for the same region of under-sequenced two technical replicates for cohesin GM12878 HiChIP data. (B) Mean profile heatmaps.

### cLoops application to Trac-looping data

We further demonstrated the generality of cLoops for calling accurate loops using the recently published Trac-looping data (Lai et al. 2018). The advantages of cLoops over the Trac-looping-methods are shown by the following: 1) Globally, loops called by cLoops were more enriched for the Trac-looping PETs compared to nearby regions (**Figure 7A**). 2) cLoops detected much more distant loops (**Figure 7B**). 3) The loops uniquely detected by cLoops were much more enriched for interacting signals (**Figure 7C**) and most of the uniquely detected loops of cLoops are more distant (**Figure 7D**). A randomly selected example shows 3 distant loops uniquely detected by cLoops as linking the significant interactions between promoters while the Trac-looping-methods detected a very close loop nearby (**Figure 7E**). 4) Hi-C signals on the Trac-looping loops also show higher enrichment of cLoops called loops compared to the Trac-looping-methods (**Figure 7F**). Even though there was no peak-calling step in cLoops, the PETs density on cLoops called loop anchors were as high as that of the Trac-looping-methods (**Figure 7G**).

**Figure 7.**
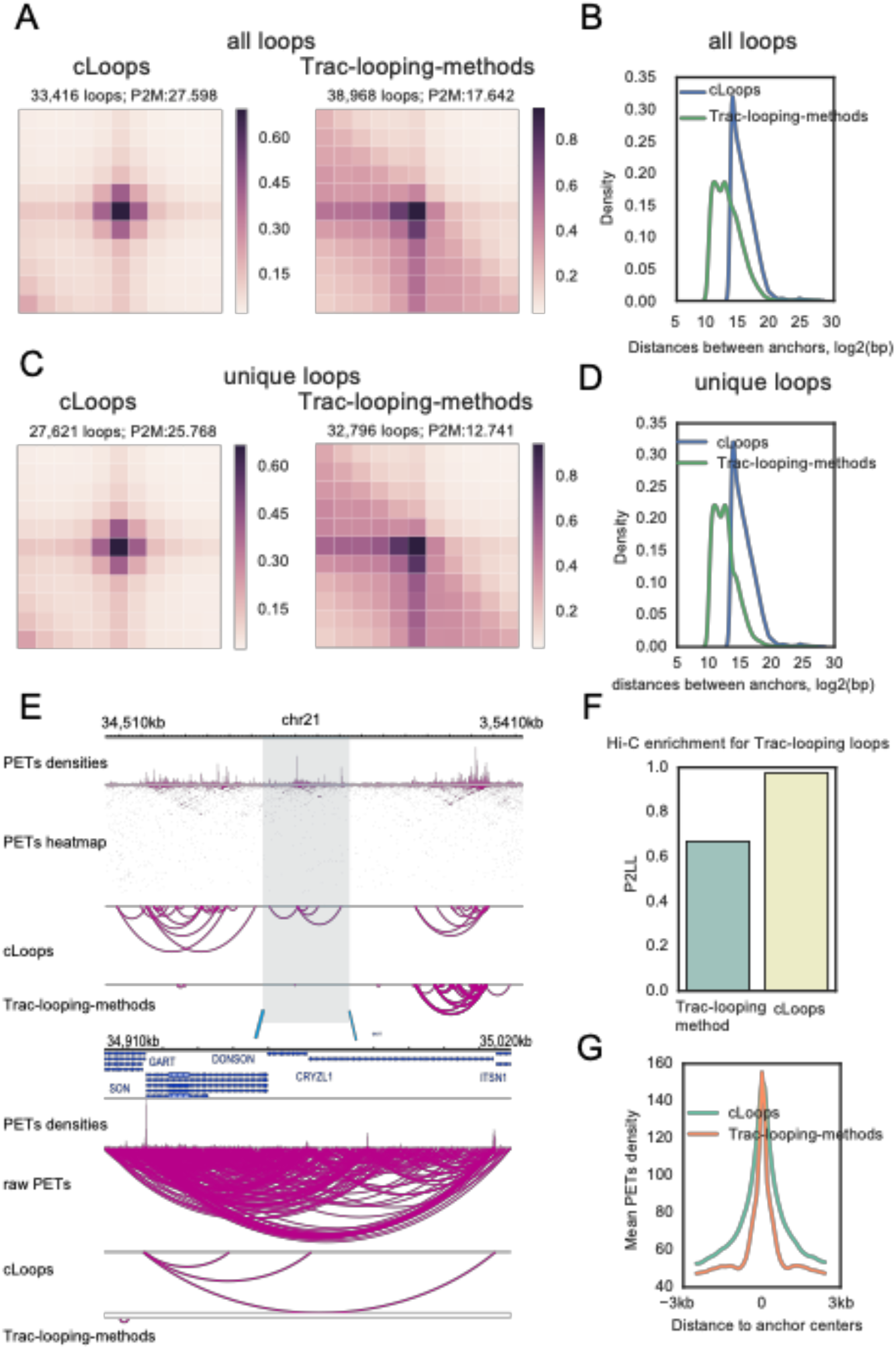
cLoops applied to Trac-looping data compared to the Trac-looping-methods. (A) Mean profile heatmaps of all loops called cLoops and the Trac-looping-method. Mapped PETs of Trac-looping data for the resting CD4 cell in BEDPE files and the Trac-looping-methods called were obtained from GSE87253. (B) Distribution of distances between loop anchors for all loops. (C) Mean profile heatmaps of unique loops called by cLoops and the Trac-looping-methods. (D) Distribution of distances between loop anchors for unique loops. (E) Randomly selected examples for cLoops and Trac-looping-methods called loops. (F) APA for evaluating the qualities of loops called from Trac-looping data using Hi-C data. The P2LL (Peak to Lower Left, suggested by Juicer) was used to show enrichment of Hi-C signal on Trac-looping loop regions. (G) Mean Trac-looping PETs densities on loop anchors.

## Discussion and conclusion

In summary we report cLoops as a new loop-calling tool based on an improved clustering algorithm, cDBSCAN, and permuted local background. We first showed the cDBSCAN clustering algorithm drastically improved speed on both simulated data and real CTCF ChIA-PET data compared to the original DBSCAN algorithm. cLoops determines the significance of loop calling by a permuted, instead of model-based, local background. These two features make cLoops applicable to ChIA-PET, HiChIP, Hi-C and Trac-looping data, other 3C-based chromatin interaction data, and yet-to-be-developed 3D mapping technologies.

## Methods

### kDBSCAN and the simulation data for performance comparison with cDBSCAN

kDBSCAN is implemented in DBSCAN with KD-tree for neighbor search. We used scipy.spatial.cKDTree for the KD-tree, which was coded in C. Specifically, the points in the 2D spaces were first built as a KD-tree, when querying for neighbors in expand cluster functions, the kdtree.query_ball_point was called. The code of cDBSCAN and kDBSCAN have been deposited to GitHub and are available at: https://github.com/YaqiangCao/cLoops_supplementaryData/tree/master/SupplementaryData/benchmarking/1.simulatedData, with file names of cDBSCAN.py and kDBSCAN.py.

100 clusters were generated randomly with centers (*X*_i_, *Y*_i_) for i from 1 to 100, where *X*_i_ and *Y*_i_ are random integers uniformly selected from (-5000, 5000). And the signal points were generated by sklearn.datasets.samples_generator.make_blobs around the centers with samples set to 10,000 and std to 0.2. The noise points were generated randomly in the space as floats. For the comparison, parameters *eps* = 0.2 and *minPts* = 5 were used. The code for benchmarking is available at: https://github.com/YaqiangCao/cLoops_supplementaryData/tree/master/SupplementaryData/benchmarking

### Mathematical model for loop significance determination

Using the permuted local background (PLB), an enrichment score is calculated as:

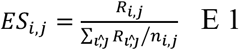

where *R_i,j_* is the number of PETs linking the anchors *i, j, n_i,j_* is the number of permutated regions, *R_l,̂j_* is the number of PETs linking the permuted regions.

FDR is defined as the ratio of permuted local regions that have more PETs than the candidate loop.

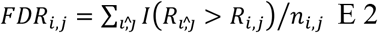

The hypergeometric test is carried out as,

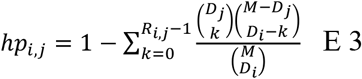

Where *M* is the total PET number in the chromosome for the region *i, j, D*_*j*_ are the total PETs in the region j (one anchor).

Using the PLB, the Poisson test can be carried out (as all numbers are integers) as,

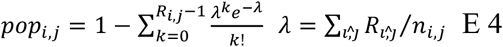

and the binomial test as,

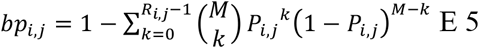

Where *P;_i,j_* is the possibility of observing 1 PETs link of the two regions normalized by the depth of the interacting regions, estimated as:

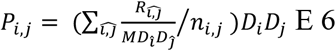

The final p-values for the hypergeometric test, Poisson test and binomial test are reported before and after correcting for multiple hypothesis testing using the Bonferroni method, multiplying the numbers of tests in that chromosome.

### Public data used

Used datasets were summarized in **Supplemental Table 1.**

### Pre-processing of ChIA-PET, Hi-C, HiChIP and Trac-looping data

The raw FASTQ files of ChIA-PET data were pre-processed by Mango (Phanstiel et al. 2015) into mapped de-duplicated intra-chromosomal PETs. To call loops with ChIA-PET2 (Li et al. 2016), ChIA-PET2 pipeline was used. Only intra-chromosomal PETs in chr1-22 and chrX were used to call loops to avoid bugs in Mango.

The raw FASTQ files of cohesin HiChIP data were first processed into mapped de-duplicated intra-chromosomal PETs by HiCUP (v0.5.4) (Wingett et al. 2015), using the genome version of hg38. HiC-Pro (v2.10.0) (Servant et al. 2015) was used to produce a fair pre-processing of data to compare performances between Hi-C loop calling tools for GM12878, K562 Hi-C data and two cohesin HiChIP technical replicates. Only intra-chromosomal PETs in chr1-22, and chrX were used to call loops. The BEDPE files for mapped resting CD4+ Trac-looping data to hg19 were obtained from GEO and replicates were combined.

### Summary of loop-calling tools comparison methods

Software version, references, key parameters, and input of loop calling tools for ChIA-PET, Hi-C and HiChIP to compare with cLoops are summarized in **Supplemental Table 2**. The Trac-looping-methods called loops for resting CD4+ (GSE87254_DHS1K_rest_3PETs_fdr.txt.gz) were obtained from GEO.

To run HiCCUPS, HIC files were generated by Juicer with the input converted from HiC-Pro’s output file with suffix of allValidPairs. When running HiCCUPS for the K562 Hi-C data, there was an error of jcuda.CudaException halt the result, so we separated K562 Hi-C data by chromosomes and run HiCCUPS for every chromosomes then combined all the result. To run HOMER, HiC-Pro’s output file with suffix of allValidPairs was converted to the required HiCSummary format. There was no specific setting or errors when analyzing Hi-C and HiChIP data by HOMER. To run Fit-Hi-C, Hi-C and HiChIP data were separated by chromosome. The output from HiC-Pro was converted to Fit-Hi-C input by HiC-Pro’s script named hicpro2fithic.py. Even for the 10kb resolution contact matrix fed to Fit-Hi-C, chr1 needs more than 75G RAM and chr2 needs about 80G. To run GOTHiC, PETs BAM file output by HiC-Pro was separated by chromosome and then feed to GOTHiC. To run diffHic, PETs BAM file output by HiC-Pro was used as input. If a specific resolution of contact matrix was needed by a tool, it was set to 10kb.

For all data tested, cLoops needs no more than 30G RAM when using one CPU. Loops called by cLoops for ChIA-PET data and the commands to run cLoops are available at: https://github.com/YaqiangCao/cLoops_supplementaryData/tree/master/SupplementaryData/loops/ChIA-PET. Loops called by cLoops for Hi-C data and the commands to run cLoops are available at: https://github.com/YaqiangCao/cLoops_supplementaryData/tree/master/SupplementaryData/loops/Hi-C. Loops called by cLoops for HiChIP data and the commands to run cLoops are available at: https://github.com/YaqiangCao/cLoops_supplementaryData/tree/master/SupplementaryData/loops/HiChIP. Loops called by cLoops for Trac-looping data and the commands to run cLoops are available at: https://github.com/YaqiangCao/cLoops_supplementaryData/tree/master/SupplementaryData/loops/Trac-looping.

### Cumulative aggregate peak analysis

The cumulative aggregate peak analysis (CAPA) was carried out according to Mango (Phanstiel et al. 2015) to evaluate loops quality called from ChIA-PET data using Hi-C. Briefly, to generate CAPA plots, we ranked loops by p-values (or FDR of ChIA-PET, hypergeometric test p-values were used for cLoops called loops) and calculated a recommend P2LL aggregate peaks analysis score by the command of APA in Juicer (Durand et al. 2016b), in a cumulative process adding 100 for most ChIA-PET loops and 20 for loops fewer than 1000 loops at a time.

### Aggregate peak analysis for loops comparison

To show the enrichment of global mean profiles of all called loops with their nearby regions for the Hi-C and HiChIP data, Juicer APA (Durand et al. 2016b) with following parameters: -n 0 –w 5 –r 5000 –u was used to get the view of centerNormedAPA and the P2M score (indicating the enrichment of loops compared to nearby regions) was used. Here –n 0 was used to analyze all loops without filtering out loops that are close to the diagonal line of the input contact matrix. For ChIA-PET data, -n was set to 0, 10, 20 and 30 (default), respectively for comparison. When carrying APA for Trac-looping loops, 1kb resolution was used both for Trac-looping and Hi-C data. The centerNormedAPA heatmaps output by Juicer APA were used for visualization comparison in genome-wide way. In a centerNormedAPA heatmap, loops are normalized at the heatmap center and indicating the loops enrichment comparing to nearby regions. If there are higher interacting regions than the center in a heatmap, may indicate there are shifts of loops boundaries or global loops quality are not good. We obtained the centerNormedAPA matrix in Juicer APA’s output of gw/enhancement.txt. We mainly used the P2M score as a loops global quality indicator due to following reasons. 1) According to Juicer (Durand et al. 2016b) documentation, the definition of P2M (Peak to Mean) score is the ratio of the central pixel to the mean of the remaining pixels, which tends to indicate the enrichment of interactions in loop regions against nearby background. We also compared the P2LL (Peak to Lower Left) score (P2LL score was suggested by Juicer APA guide) and its related ZscoreLL, which is the ratio of the central pixel to the mean of the mean of pixels in the lower left corner. 2) If there are too many loops fed to Juicer APA, for example 748,786 GM12878 Hi-C loops output by GOTHiC, Juicer APA will crash that leads to no output of gw/measures.txt file, which records the global P2LL score and other indicators. Meanwhile, there is always a file named “enhancement.txt” recording P2M score for every loop when feeding loops to Juicer APA one chromosome at a time.

### Visualization of example loops

Juicebox (Durand et al. 2016a) was used to show loops with a resolution for the heatmap of 5kb.

## AUTHOR CONTRIBUTIONS

YQC and JDJH designed the project; YQC and XWC implemented the cDBSCAN algorithm; YQC and DSA implemented the local permutated significance test model. All authors contributed to data analysis, interpretation and wrote the paper.

